# Different analysis methods of Scottish and English child physical activity data explain the majority of the difference between the national prevalence estimates

**DOI:** 10.1101/408179

**Authors:** Chloë Williamson, Paul Kelly, Tessa Strain

**Affiliations:** Physical Activity for Health Research Centre, Institute of Sport, Physical Education, and Health Sciences, Moray House School of Education and Sport, University of Edinburgh; MRC Epidemiology Unit, University of Cambridge, Cambridge, CB2 0QQ, United Kingdom

**Keywords:** children, physical activity, surveillance

## Abstract

**Background:** The percentage of children in Scotland and England meeting the aerobic physical activity recommendation differ greatly according to the estimates derived from the respective national health surveys. The Scottish Health Survey (SHeS) usually estimates that over 70% meet the guidelines; Health Survey for England (HSE) estimates are usually below 25%. It is plausible that these differences originate from different analysis methods. The HSE monitor the percentage of children that undertake 60 minutes of moderate-to-vigorous physical activity on each day of the week (‘the daily minimum method’ (DMM)). The SHeS monitor the proportion that undertake at least seven sessions of moderate-to-vigorous physical activity, with an average daily duration over 60 minutes (‘the average method’ (WAM)). We aimed to establish how great an influence this difference in analysis methods has on the prevalence estimates.

**Methods:** Physical activity data from 5-15 year olds in the 2015 HSE and SHeS were reanalysed (weighted n=3840 and 965, respectively). Two comparable pairs of estimates were derived: a DMM and WAM estimate from the HSE not including travel to/from school, and WAM estimates from the HSE and the SHeS including travel to/from school. It is not possible to calculate a DMM estimate from the SHeS due to the way the questions are asked. Results were presented for the total samples, and by sex and age sub-groups.

**Results:** The HSE WAM estimate was 31.7 (95% CI: 30.2-33.3) percentage points higher than the DMM estimate (54.3% (95% CI: 52.6-56.0) and 22.6% (95% CI: 21.2-24.1) respectively). The magnitude of this difference differed by age group but not sex. When comparable WAM estimates were derived from the SHeS and the HSE, the SHeS was 11.8 percentage points higher (73.6% (95% CI: 69.8-77.1) and 61.8% (95% CI: 60.2-63.5) respectively). The magnitude of this difference differed by age group and sex.

**Conclusions:** The results indicate that the difference in the analysis method explains the majority (approximately 30 percentage points) of the difference in the child physical activity prevalence estimates between Scotland and England. These results will help those involved in national surveillance to determine how to increase comparability between the U.K. home nations.

## Background

There are a number of surveys within the UK home nations that estimate the proportion of children meeting the aerobic physical activity (PA) recommendation (≥60 minutes of moderate-to-vigorous PA (MVPA) every day [1]). For example, the national health surveys and the Health Behaviour in School-Aged Children survey (HBSC) [2-4]. Cross-country comparisons are inevitable, indeed often encouraged [5]. The study designs, measurement instruments, methods of administration, and age ranges included vary between surveys, but most estimate between 10-30% of children in their target population meet the 60 minute threshold [2-4,6,7]. The Scottish Health Survey (SHeS) is an exception to this, having reported estimates of over 70% annually since 2008 [8]. The conclusion that Scottish children are more active than those in other home nations is not supported when the Scottish, English and Welsh HBSC survey results are compared [4-6,9]. The comparable age and sex sub-group prevalence estimates from the different nations’ surveys are all within six percentage points of each other, ranging between 11-30% [4, 6,9].

A plausible contributing factor is that the SHeS estimates are for 2-15 year olds, while most others are for older age groups (e.g. 5-15 year olds in Health Survey for England (HSE), or 11-16 year olds in the HBSC surveys). This may inflate the SHeS estimates because the youngest are the most active age group, although the evidence suggests it would account for a few percentage points at most [8]. There is a separate issue around whether 2-4 year olds should be included in the prevalence estimate given that the recommendation in question applies to 5-18 years olds [1]. Those under 5 years that can walk unaided are recommended to be physically active for at least 3 hours per day [1]. The reason for their inclusion in the SHeS estimates is to maintain the trend data that began before a distinction was made between the age groups.

Some have attributed the high prevalence figures to the SHeS questionnaire over-estimating MVPA time [10]. This line of thought is based on a convergent validity study comparing the SHeS questionnaire^1^ to uniaxial waist-worn accelerometers in 130 6-7 year olds [11]. The estimates for total daily MVPA derived from the questionnaire were approximately two hours higher than those from accelerometry. As the SHeS does not specifically explain to respondents that they should only report activity of at least moderate intensity [12], it is easy to see how these conclusions are drawn.

However, the overwhelming majority of activities that are prompted are considered by the Youth Compendium of Physical Activities would consider to be of at least moderate intensity (e.g. sweeping leaves, running about, football; [13]). Also, even if there were some light intensity activities reported, one would expect there to be a comparable degree of reporting in the HSE where the same types of activities are prompted, just under different categories and in a slightly different order (see **Table 1**) [14]. This does not appear to be the case as there was still a difference of over 50 percentage points between the 2015 estimates (73% in SHeS and 22% in HSE) [2, 3].

**Table 1.** The measurement of child physical activities in the 2015 Scottish Health Survey and 2015 Health Survey for England.

| Specific physical activity | SHeS 2015 category of activity | HSE 2015 category of activity | Included in estimates presented in this study |  |  |  |
| --- | --- | --- | --- | --- | --- | --- |
|  |  |  | (1) HSE DMM no School Travel estimate<sup>a</sup> | (2) HSE WAM no School Travel estimate | (3) HSE WAM All Activity estimate | (4) SHeS WAM estimate |
| Activity as part of school lessons | School based activity | School lessons | ✓ | ✓ | ✓ | ✓ |
| Walking to school | Walking | Travel to school<sup>b</sup> | X<sup>b</sup> | X | ✓ | ✓ |
| Cycling to school | Sports and exercise activities |  | X<sup>b</sup> | X | ✓ | ✓ |
| Walking (not to school) | Walking | Informal activities | ✓ | ✓ | ✓ | ✓ |
| Active play activities | Active play |  | ✓ | ✓ | ✓ | ✓ |
| Heavy housework/gardening | Housework<sup>c</sup> |  | ✓ | ✓ | ✓ | ✓<sup>c</sup> |
| Sport and exercise activities | Sport and exercise activities | Formal activities | ✓ | ✓ | ✓ | ✓ |
| Any other activities similar to those prompted under active play and/or sport and exercise activities | Active play | Other informal/formal activities | ✓ | ✓ | ✓ | ✓ |

SHeS – Scottish Health Survey, HSE – Health Survey for England, WAM – Weekly Average Method, DMM – Daily Minimum Method (see article text for descriptions). a: any minor differences between this estimate and the published headline HSE figures because those not at school are allocated 0 for school lesson activity rather than being excluded; b: the HSE does not ask the specific days (only number of days) on which the child either walked or cycled to, or part of the way to, school/nursery/playgroup. This means it cannot be included in any prevalence estimate using the DMM; c: only asked of those aged 8-15 years.

There is a separate but important debate about whether it is appropriate to measure compliance to the child MVPA recommendation via accelerometers. This is beyond the scope of this paper, but readers are referred to RP Troiano, JJ McClain, RJ Brychta and KY Chen [15] who state that ‘evaluation of PA guideline adherence based on accelerometer outcomes is inappropriate because the behavioural metrics used to develop the guidelines differ conceptually from device-based measures of MVPA.’

Given that the SHeS and HSE questionnaires are so similar, yet produce such different prevalence estimates, we investigated how the data were processed. A subtle but potentially important difference between the SHeS and HSE surveys is how the frequency of activities are reported, which has implications for how the data are processed. In the SHeS, children or their parents are asked to report how many sessions of each activity they have done in the previous seven days [12]. In the HSE, activities are reported for specific days in the previous seven [14]. It is therefore not possible to ascertain whether child respondents to the SHeS achieve ≥60 minutes of MVPA on each specific day. Instead, one can calculate whether a child has reported ≥7 sessions, and whether the total weekly duration is equivalent to an average ≥60 minutes per day. For the rest of this paper, we use the term ‘Weekly Average Method’ (WAM) to describe this approach. This differs from the ‘Daily Minimum Method’ (DMM) used by the HSE where a child is judged to meet the recommendation if they report ≥60 minutes per day on each specific day of the preceding week (see **Figure 1**).

**Figure 1.**
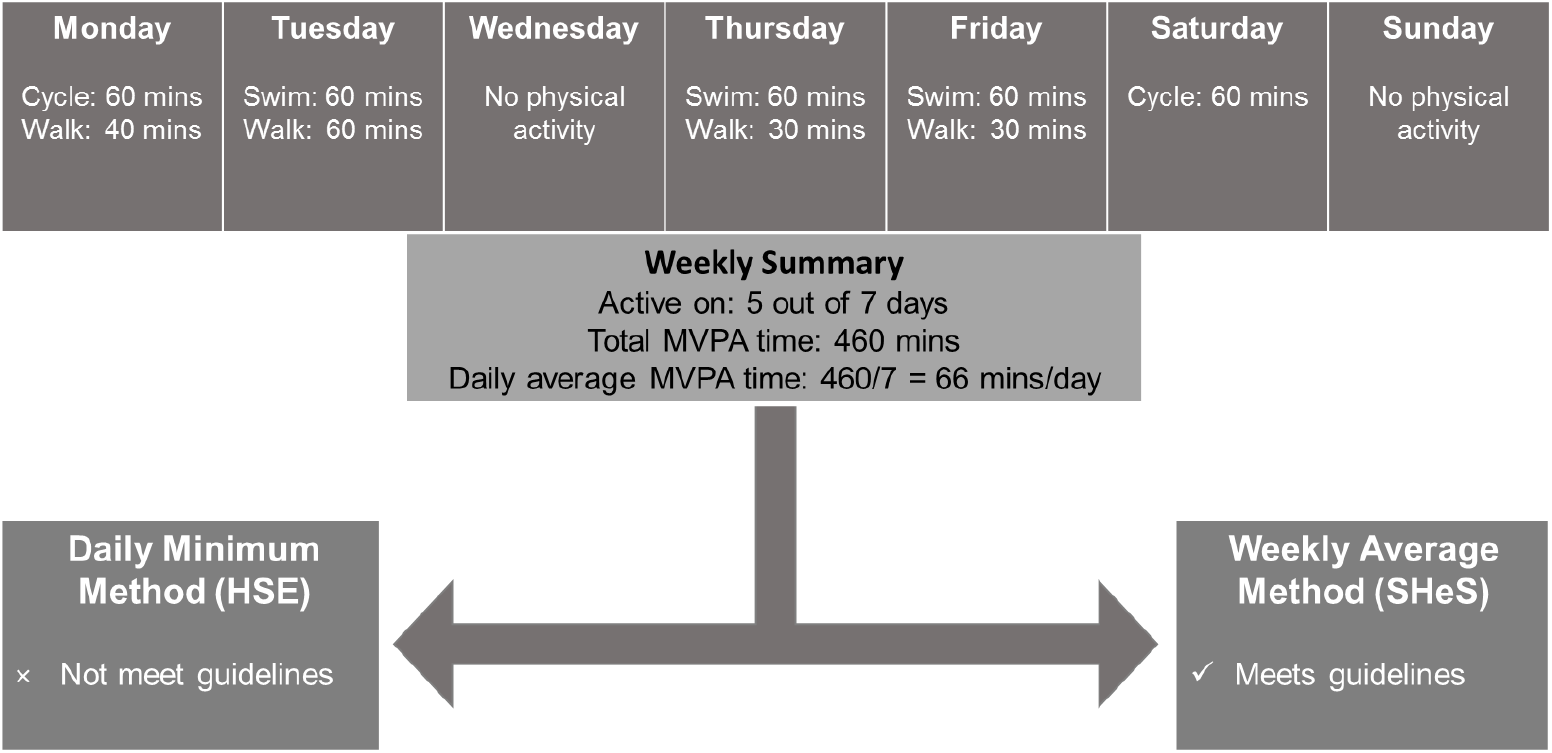
Demonstration of how the use of ‘Daily Minimum Method’ or the ‘Weekly Average Method’ may lead to different results from the same data.

We hypothesise that it is this different analysis method that is the key driver behind the differences in prevalence estimates in the SHeS and the HSE. This is important to understand because it will offer strategies for appropriate changes to be made to surveillance methods to increase comparability between home nations. It is timely as such changes may be precipitated by the 2018 update of the CMO PA Recommendations [16]. We will test this hypothesis by reanalysing HSE and SHeS data to generate comparable estimates between Scotland and England. **Figure 2** provides an overview of what methods (WAM or DMM) are possible in the two surveys, and what comparisons can be drawn.

**Figure 2.**
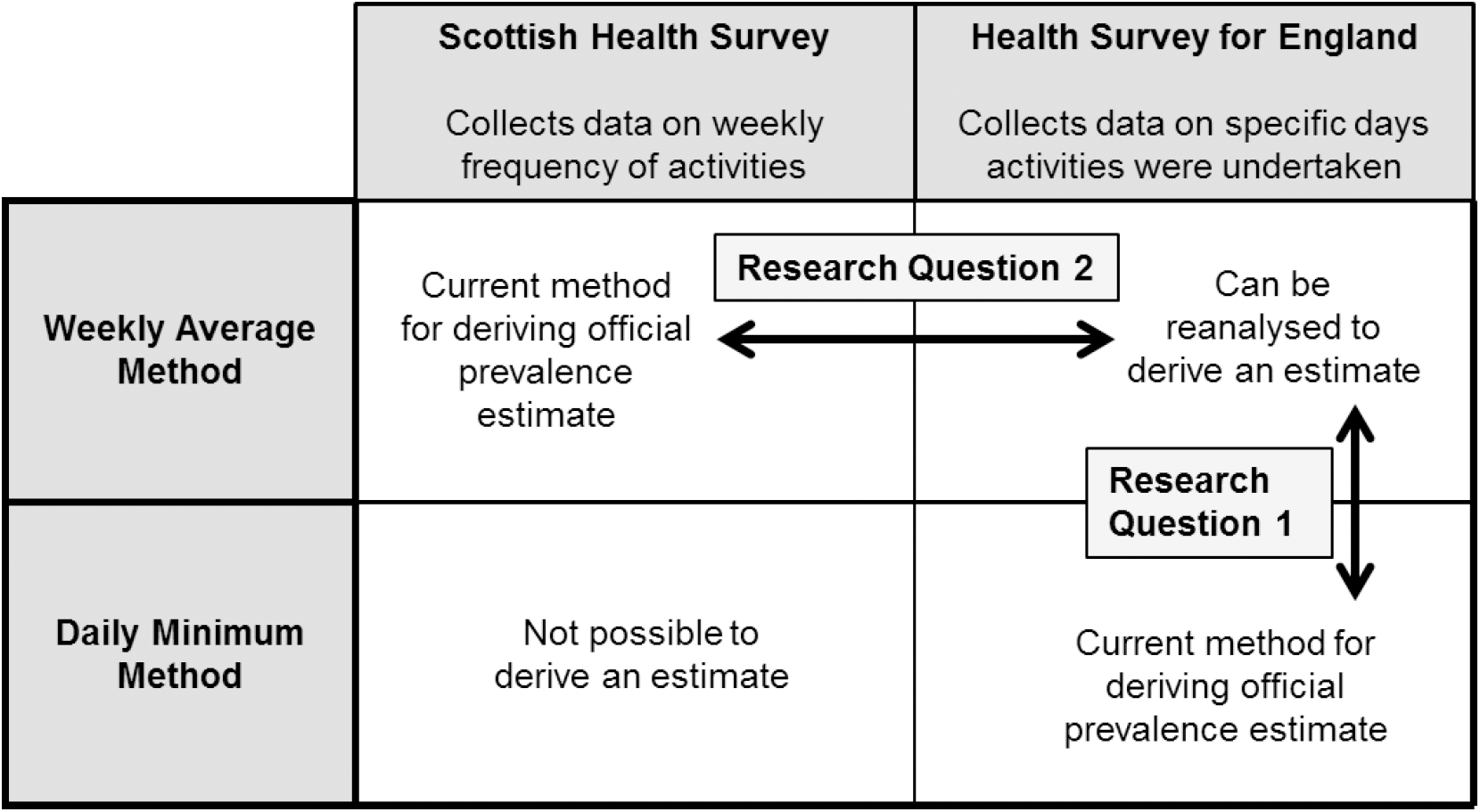
Schematic overview of what analysis methods are possible in the Scottish Health Survey and Health Survey for England, and the research questions of the study.

Our two research questions are: (1) what is the percentage point difference between comparable WAM and DMM estimates for England? (2) What is the percentage point difference between comparable WAM estimates for Scotland and England? These differences will be assessed for all children, and by age and sex sub-groups.

## Methods

### Data Sources

The study was approved by the Moray House School of Education ethics committee and all authors with access to the data agreed to the UK Data Archive End User Licence. The 2015 HSE and SHeS datasets were downloaded from the UK Data Archive on 6^th^ April 2018 [17, 18], previously preliminary work was undertaken on the respective 2012 datasets. Both surveys are sampled so that they are nationally representative of their respective national populations that live in private households after weighting on key demographic characteristics; further details are in the surveys’ technical reports [12, 14].

### Sample

The sample age range was restricted to 5-15 year olds. Those younger have different PA guidelines [1]. Those older for whom the recommendation does apply (16-18 year olds) are treated as adults by both surveys and so responded to a different questionnaire. Those responding ‘don’t know’ or who refused to answer a question were excluded (<2%). Those not at school were included in all analyses but were allocated zero time/frequency for any activity at or involved in the travel to/from school. This left 4096 respondents to the 2015 HSE and 946 respondents to the 2015 SHeS with complete PA data.

### Physical activity measurement

**Table 1** shows that the HSE and SHeS ask about the same activities but that they split under different categories. This means the ordering is slightly different. Both surveys have a recall period of the seven days prior to interview. The full questionnaire transcripts are published in the technical reports [12, 14]. Parents answered on behalf of children aged 5-12, and children aged 13-15 answered themselves.

Four summary measures of compliance to the PA recommendation were derived (see **Table 1**):

1. from the 2015 HSE using the DMM. This did not include travel to/from school because data were reported as a weekly frequency rather than on specific days.
2. From the 2015 HSE using the WAM, also excluding travel to/from school such that it was comparable with (1).
3. From the 2015 HSE using the WAM, including travel to/from school as a weekly frequency was not a limitation for this method.
4. From the 2015 SHeS using the WAM, including all activity so it was comparable with (3).

The mean differences between the pairs of estimates (2)-(1) (research question 1 on **Figure 2**) and (4)-(3) (research question 2 on **Figure 2**) were calculated. The first comparison indicated the magnitude of the difference that the analysis method makes, because it uses the two methods in the same sample. The second comparison was between the WAM estimates from the two different surveys. This was an indication of how much of the difference was not explained by the analysis method (possibly true difference, or other methodological differences that could not be standardised).

Statistical differences by sex and age group (5-7, 8-10, 11-12, 13-15 years) were assessed using adjusted Wald F-tests. All estimates were weighted to account for non-response and selection bias. Analyses were carried out using Stata/SE v14.2, Texas, USA.

## Results

The weighted sample sizes were n=3840 and n=947, for the 2015 HSE and SHeS respectively (see **Table 2**). The breakdown of the samples by sex and age group were near identical (<2 percentage point differences).

**Table 2.** Descriptive characteristics of the analysis samples from each survey.

|  | **HSE 2015 sample** |  | **SHeS 2015 sample** |  |
| --- | --- | --- | --- | --- |
|  | **weighted n** | **%** | **weighted n** | **%** |
| **Total sample** | 3840 | 100 | 965 | 100 |
| **Boys** | 1958 | 51.0 | 493 | 51.1 |
| **Girls** | 1881 | 49.0 | 472 | 48.9 |
| **5-7 years** | 1133 | 29.5 | 281 | 29.1 |
| **8-10 years** | 1058 | 27.5 | 272 | 28.2 |
| **11-12 years** | 723 | 18.8 | 167 | 17.4 |
| **13-15 years** | 927 | 24.1 | 244 | 25.3 |

When the 2015 HSE sample was analysed according to the DMM and the WAM (both not including travel to/from school), the WAM estimate was 31.7 percentage points higher than the DMM estimate (22.6% (95% CI: 21.2-24.1) and 54.3% (95% CI: 52.6-56.0); see **Figure 3, Table 3**). An increase of comparable magnitude was evident in both boys and girls (p>0.05 for difference between sexes), but differed by age group (p<0.05). The differences between the estimates for the youngest two age groups (5-7 year olds and 8-10 year olds) were within two percentage points of the sample mean. The difference was highest for 11-12 year olds (36.6 percentage points (95% CI: 32.7-40.6)) and lowest for 13-15 year olds (28.5 (95% CI: 25.5-31.6).

**Table 3.** The mean percentage point differences and 95% confidence intervals between comparable estimates.

|  | (1) HSE - DMM (no travel to/from school) | (2) HSE - WAM (no travel to/from school) | Mean difference: (2) HSE WAM – (1) HSE DMM | (3) HSE – WAM (including travel to/from school) | (4) SHeS - WAM | Mean difference: (4) SHeS WAM – (3) HSE DMM |
| --- | --- | --- | --- | --- | --- | --- |
|  | % (95% CI) | % (95% CI) | Percentage point (95% CI) | % (95% CI) | % (95% CI) | Percentage point (95% CI) |
| **Total sample** | 22.6 (21.2, 24.1) | 54.3 (52.6, 56.0) | 31.7 (30.2, 33.3) | 61.8 (60.2, 63.5) | 73.6 (69.8, 77.1) | 11.8 (8.3, 15.3) |
| **Boys** | 24.8 (22.8, 27.0) | 57.6 (55.2, 60.0) | 32.8 (30.6, 35.0) | 64.5 (62.2, 66.8) | 79.6 (75.0, 83.5) | 15.0 (10.6, 19.4) |
| **Girls** | 20.3 (18.4, 22.3) | 50.9 (48.5, 53.3) | 30.6 (28.4, 32.8) | 59.1 (56.7, 61.4) | 67.4 (61.3, 73.0) | 8.3 (3.0, 13.7) |
|  |  |  | *p>0.05* |  |  | *p<0.05* |
| **5-7 years** | 29.1 (26.4, 32.0) | 59.4 (56.3, 62.3) | 30.3 (27.5, 33.0) | 64.6 (61.6, 67.4) | 77.8 (71.8, 82.9) | 13.2 (7.4, 19.1) |
| **8-10 years** | 26.7 (23.9, 29.6) | 59.4 (56.3, 62.5) | 32.8 (29.7, 35.8) | 65.4 (62.3, 68.4) | 83.1 (77.7, 87.5) | 17.7 (12.1, 23.3) |
| **11-12 years** | 18.4 (15.3, 22.0) | 55.0 (50.8, 59.1) | 36.6 (32.7, 40.6) | 65.2 (61.1, 69.1) | 69.2 (59.6, 77.3) | 4.0 (-5.5, 13.4) |
| **13-15 years** | 13.3 (11.2, 15.8) | 41.9 (38.6, 45.2) | 28.5 (25.5, 31.6) | 51.9 (48.5, 55.3) | 60.9 (53.0, 68.2) | 9.0 (1.3, 16.6) |
|  |  |  | *p<0.05* |  |  | *p<0.05* |

HSE: Health Survey for England; SHeS: Scottish Health Survey; WAM – Weekly Average Method, DMM – Daily Minimum Method (see article text for descriptions). P-values refer to the testing of the null hypotheses that the proportion is equal across sex and age sub-groups.

**Figure 3.**
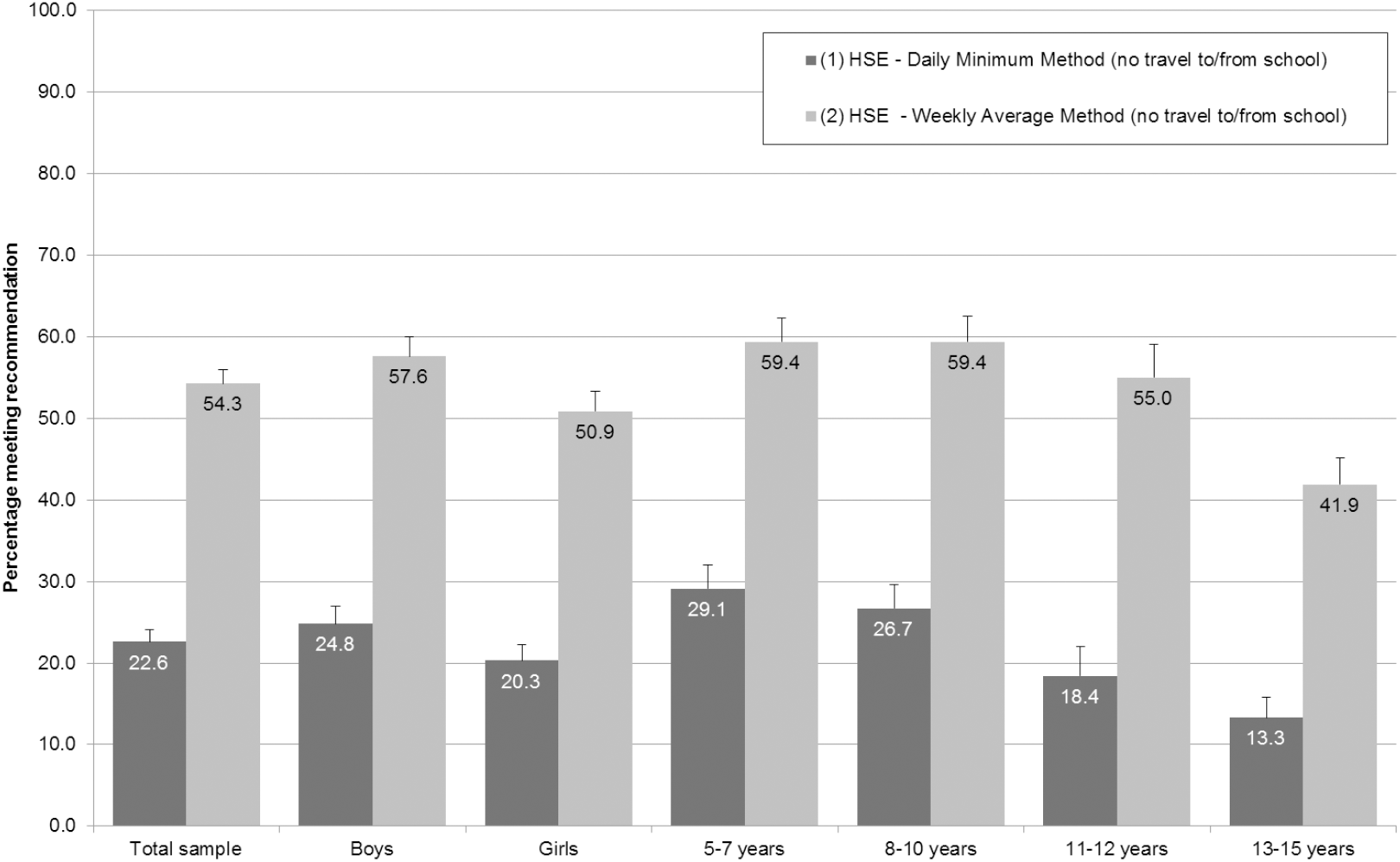
The percentages of children in the 2015 Health Survey for England meeting the aerobic physical activity recommendation according to the analysis methods.

When the comparable estimates (i.e. including all reported activities) were derived from the 2015 HSE and the 2015 SHeS using the WAM, there was an 11.8 percentage point (95% CI: 8.3-15.3) difference (61.8% (95% CI: 60.2-63.5) for the HSE and 73.6% (95% CI: 69.8-77.1) for the SHeS; see **Figure 4, Table 3**). This difference was greater amongst boys than girls (15.0 percentage points (95% CI: 10.6-19.4) and 8.3 (95% CI: 3.0-13.7) respectively; p<0.05). The difference was highest amongst 8-10 year olds (17.7 percentage points (95% CI: 12.1-23.3)) and lowest amongst 11-12 year olds (4.0 (95% CI: −5.5-13.4); p<0.05 for difference between age groups).

**Figure 4.**
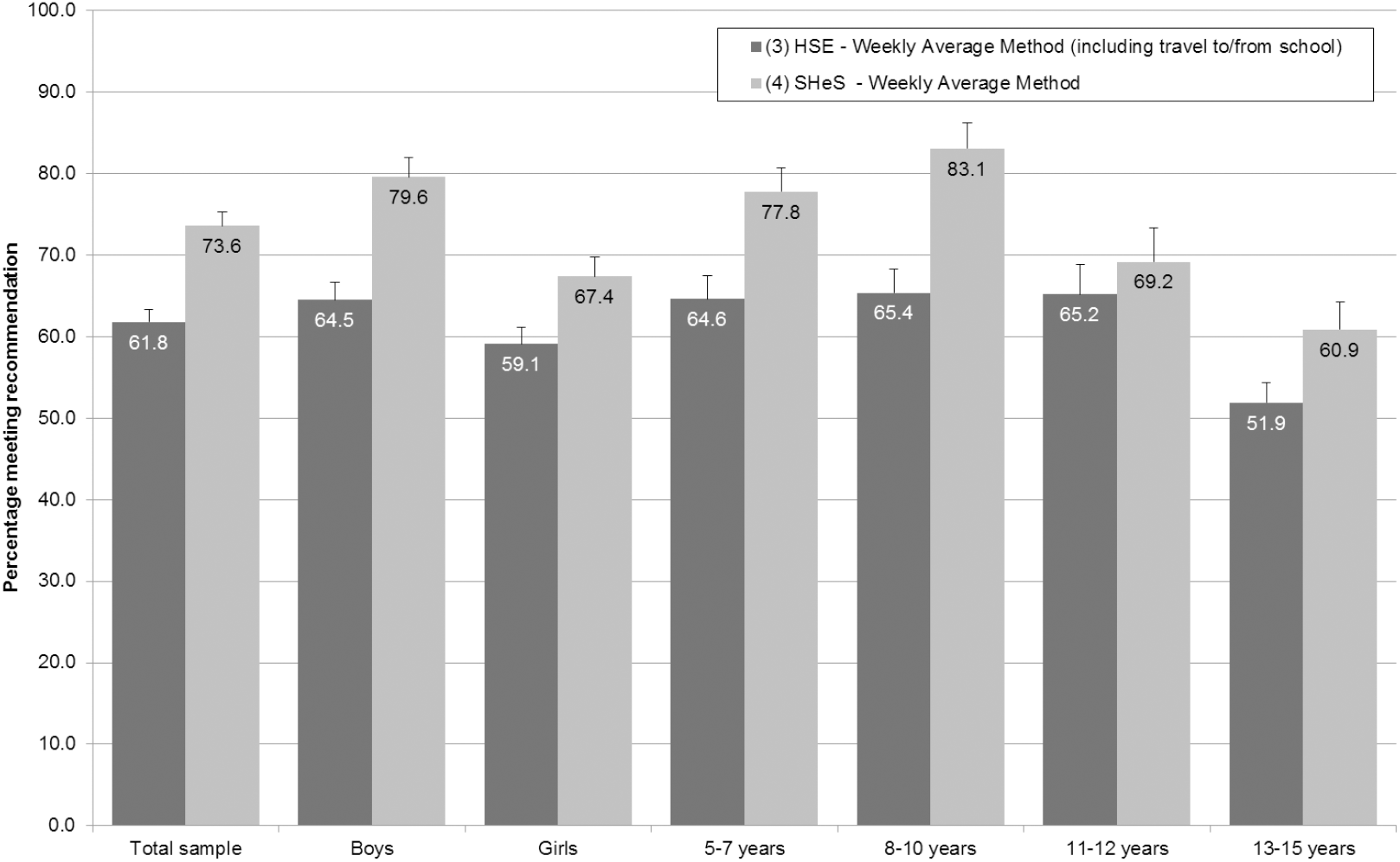
The percentages of children in the 2015 Health Survey for England and the 2015 Scottish Health Survey meeting the aerobic physical activity recommendation according to the different analysis methods.

## Discussion

These findings show that small differences in the analysis method can lead to a critical difference in prevalence. Our results indicate the majority of the difference between the SHeS prevalence estimate and those from other UK national surveys is due to a different interpretation of the’ 60 minutes per day’ recommendation. This is shown by the 31.7 percentage point increase (95% CI: 30.2-33.3) in prevalence when the HSE sample was reanalysed according to the WAM rather than the DMM. There remains, however, an 11.8 percentage point (95% CI: 8.3-15.3) difference between the comparable WAM estimates from the SHeS and the HSE. This may in part be due to a true difference between the PA levels of children in Scotland and England, but may also be due the different categorisation of activities and ordering of questions in the SHeS and the HSE.

This leads to an important conclusion; small differences in analysis methods could invalidate cross-country comparisons and at worst will lead to misguided policy and practice. This highlights the need for caution when interpreting the results of cross-country comparison projects such as the Active Healthy Kids Report Cards and the GoPA country cards [5, 19]. While they may serve a political or advocacy purpose, any interpretation that county Y has a higher/lower PA prevalence than country Z should be approached with caution. We should question: how were data collected and analysed, before we ask: are prevalence estimates real and why do they exist?

This naturally leads to the question: what is the most appropriate interpretation of the guideline? The Technical Report accompanying the 2011 UK PA guidelines is clear that they intended the DMM interpretation [20]. The recommendation for daily physical activity was included to ‘promote a pattern of regular PA’ and to assist communication of a simple message. However, they also state that ‘there is no evidence that missing one day (or two or three days) in a week has any measurable negative effect on health’ [20]. However, there is room for discussion over the evidence used to support this decision. It may be that a “rest day” each week allows for better recovery, rest and physiological adaption if sufficient activity completed over 7 day period. Importantly it may also be better for affect and enjoyment to take a break; the potential impact on long term habit formation is clear.

In light of this, we call on developers of PA recommendations to also include guidance on how to collect, process and analyse data. This should be considered a necessary step in the implementation of any recommendations. Our results show that even small room for interpretation or “researcher or analytical degrees of freedom” can lead to huge difference in prevalence estimates. This is likely to be particularly relevant in future should device-based measures be introduced, given the permutations in processing and analysis [21, 22]. The UK PA guidelines are being reviewed this year, and, for the first time, there is a specific sub-group to consider implementation and surveillance issues. This is a useful opportunity to make sure that the surveillance methods are able to implement the guidelines as intended.

These results are comparable to a recent study that analysed accelerometry data from a Scottish child cohort study through both the WAM and DMM approaches [23]. The DMM produced an estimate of 11% whilst the WAM produced an estimate of 68%. Taken together, they show that it is important not to conflate the issues of whether device-based surveillance of child PA should be introduced in Scotland with the issues of the high prevalence estimates. Whilst there may be many other relevant considerations for changing national surveillance of child PA to accelerometry, generating comparable prevalence figures to the other national surveys within the UK could be achieved through changes to the questionnaire and/or analysis method.

The strengths of this study are that it is the first to attempt to address directly address the issue of why the prevalence figures are so different between Scotland and England. Through the research questions posed and the study design chosen, we were able to isolate the issue of the recommendation interpretation from other persisting concerns around the validity and reliability properties of the measurement instruments. Also, the large effect size increases confidence in the conclusions. Early findings of this work were shared with the Scottish Government and those running the SHeS, and have been part of considerations around changes to the measurement instrument. They will continue to do so as the UK PA guidelines are re-evaluated during 2018. The findings have also relevance for a wider audience: D Jurakic and Ž Pedišić [24] showed that differences in recommendation interpretation occur beyond the U.K‥

The main limitation of this study was that we were unable to eliminate all differences between the questionnaires, which may have affected the comparisons between the HSE and SHeS WAM estimates. It is also possible that the requirement of the HSE to report activity on specific days made it somewhat like a diary, leading respondents to answer in a different manner to those in the SHeS sample. It has been argued that diaries improve recall and reduce the potential for social desirability bias compared to general questionnaires, as stronger associations with estimates derived from doubly labelled water have been demonstrated [25]. We were also not able to account for the stratified and clustered sampling procedures when calculating the 95% CIs because of the small number of primary sampling units in some HSE strata. Therefore, the true variance on the estimates may be fractionally larger than those presented.

In conclusion, this study has demonstrated that it is the difference in analysis method that explains the majority of the difference in the child PA prevalence estimates between Scotland and England. These results will help those involved in national surveillance to determine how to increase comparability between the U.K. home nations.

## Abbreviations

HSE: Health Survey for England
SHeS: Scottish Health Survey
HBSC: Health Behaviour in School-Aged Children
MVPA: Moderate-to-vigorous physical activity
PA: Physical activity
WAM: Weekly Average Method
DMM: Daily Minimum Method

## Declarations

### Ethical approval

The study was approved by the Moray House School of Education ethics committee and all authors with access to the data agreed to the UK Data Archive End User Licence.

*Consent for publication*

Not applicable

*Availability of data and materials*

The datasets analysed during the current study are available in the UK Data Archive, at http://doi.org/10.5255/UKDA-SN-7480-1 and http://doi.org/10.5255/UKDA-SN-7417-3.

## Authors’ contributions

TS and PK developed the original research question and supervised CW who designed the study, undertook the initial analyses and drafted the manuscript. TS finalised the analyses. All authors contributed and commented on subsequent drafts.

## Acknowledgements

We would like to thank the participants of the Scottish Health Survey and the Health Survey for England, and NatCen and ScotCen Social Research for managing the surveys and processing the data.

## Footnotes

1 At the time, this was also the HSE questionnaire. The HSE questionnaire subsequently changed (in 2008) the way activities were grouped into categories and the ordering of questions. It also changed to ask on which specific day activities were undertaken, rather than a weekly frequency.

